# Dietary modulation alters susceptibility to *Listeria monocytogenes* and *Salmonella typhimurium* in a gut microbiota-independent manner

**DOI:** 10.1101/2021.06.09.447822

**Authors:** Mathis Wolter, Alex Steimle, Jacques Zimmer, Mahesh S. Desai

## Abstract

Food safety has considerably improved worldwide, yet infections with food-borne human enteric pathogens, such as *Listeria* spp. and *Salmonella* spp., still cause numerous hospitalizations and fatalities. Thus, the need to shed more light on the mechanisms of enteropathogenesis is apparent. Since dietary alterations, including fiber deficiency, might impact the colonization resistance by the gut microbiota, studying diet–microbiota–pathogen axis holds promise in further understanding the pathogenesis mechanisms. Using a gnotobiotic mouse model containing a 14-member synthetic human gut microbiota (14SM), we have previously shown that dietary fiber deprivation promotes proliferation of mucin-degrading bacteria leading to a microbiota-mediated erosion of the colonic mucus barrier, which results in an increased susceptibility towards the rodent enteric pathogen *Citrobacter rodentium.* Here, we sought to understand how low-fiber diet affects susceptibility to *Listeria monocytogenes* and *Salmonella typhimurium* infections in our 14SM gnotobiotic mouse model, in BALB/c and C57BL/6N backgrounds, respectively. Intriguingly and in contrast to our results with *C. rodentium*, we observe that depriving mice of dietary fiber protected them from infections with the pathogens compared to mice fed a standard chow. The microbiota delayed the overall pathogenicity as compared to the onset of disease observed in germ-free control mice; nevertheless, we observe the same effect of diet in germ-free mice, suggesting that the susceptibility is microbiota independent. Our study points out an important observation that dietary fiber plays a crucial role on either the host susceptibility, the virulence of these pathogens, or both, which would be judicious to design and interpret future studies.

**Importance:** Human enteric pathogens *Listeria monocytogenes* and *Salmonella typhimurium* are employed as classical models in rodent hosts to understand the pathogenesis mechanisms of food-borne pathogens. Research in the past decade has stressed importance of the composition of the gut microbiota in modulating susceptibility to these pathogens. Our results—using gnotobiotic mice and germ-free control animals—additionally suggest that the dietary fiber components dominate the impact of enteropathogenic virulence over the pathogenicity-modulating properties of the gut microbiota. The significance of our research is in the need to carefully choose a certain chow when performing the enteropathogen-associated mouse experiments and to cautiously match the rodent diets when trying to replicate experiments across different laboratories. Finally, our data underscore the importance of germ-free control animals to study these pathogens, as our findings would have been prone to misinterpretation in the absence of these controls.

## Main text

The gut microbiota confers colonization resistance against invading pathogens by nutrient competition, and by maintaining the host immune homeostasis and the mucosal barrier integrity (1). Deficiency of dietary fiber might negatively affect these host-beneficial properties of the microbiome (2). Since dietary fiber consumption in Western countries is below the recommended intake of 25–35 g per day (3), such dietary habits might contribute to the observed incidence of enteric pathogen infections in the Western world. Using a well-characterized 14-member synthetic human gut microbiome (14SM) in gnotobiotic mice, we have previously demonstrated that dietary fiber deprivation leads to an increase in the mucin-degrading gut microbiome, which erodes the colonic mucus barrier (4). We further showed that the reduced mucus barrier enhances susceptibility to infection with *Citrobacter rodentium* (4), a rodent pathogen used to model human enteropathogenic and enterohaemorrhagic *E. coli* infections (5). Since the intestinal mucus barrier is a first line of innate defense (1), here, we hypothesized that the diet-induced mucus erosion might also increase susceptibility to other enteric pathogens.

Food safety has increased considerably in recent years, yet food-borne enteric pathogens such as *Listeria* spp. and *Salmonella* spp. remain a major source of disease, even in industrialized countries (6, 7). Since dietary alterations, including fiber deficiency, might alter colonization resistance by the gut microbiota to enteric pathogens (1), understanding the interconnections in the diet–microbiota–pathogen axis might help to shed light on hitherto unexplored pathogenesis mechanisms. It has previously been shown that mice lacking the *Muc2* gene, which encodes for the major constituent glycoprotein of the colonic mucus layer, are more susceptible towards *L. monocytogenes* and *S. typhimurium* and infection (8, 9). Notably, a similar increase in susceptibility was observed for *C. rodentium* in *Muc2^−/−^* mice (10), a result that we could recapitulate in wild-type, fiber-deprived mice with the reduced mucus barrier (4). Thus, we leveraged our 14SM gnotobiotic model to investigate how dietary fiber deprivation and/or eroded mucus barrier affect the host susceptibility towards infections with the intracellular enteric pathogens *L. monocytogenes* and *S. typhimurium*.

For this purpose, we employed BALB/c and C57BL/6 mouse strains for infections with *L. monocytogenes* and *S. typhimurium,* respectively; the choice of the host strains for the respective pathogens is based on the preference of the specific pathogens for the strains, as shown by previous studies (11–14). We colonized the 6–10 weeks old, germ-free (GF) mice with the 14SM community and confirmed colonization of all 14 strains by qPCR using strain-specific primers as described previously (4, 15). For six days after colonization, the mice were kept on a standard mouse chow, which we call fiber-rich (FR) diet. After six days, half of the mice were switched to a fiber-free (FF) diet. After an additional 20 days, BALB/c mice were infected via intragastric gavage with 10^9^ colony-forming units (CFU) of *L. monocytogenes* and C57BL/6 mice were infected with 10^8^ CFU of *S. typhimurium.* Disease progression after infection was monitored for up to 10 days (**Fig. 1A, upper mouse groups**). Age- and sex-matched GF BALB/c and C57BL/6 mice were used as controls and also fed either FR- or FF-diets before being subjected to *L. monocytogenes* and *S. typhimurium* infection **(Fig. 1A, lower mouse groups)**.

**FIG 1.**
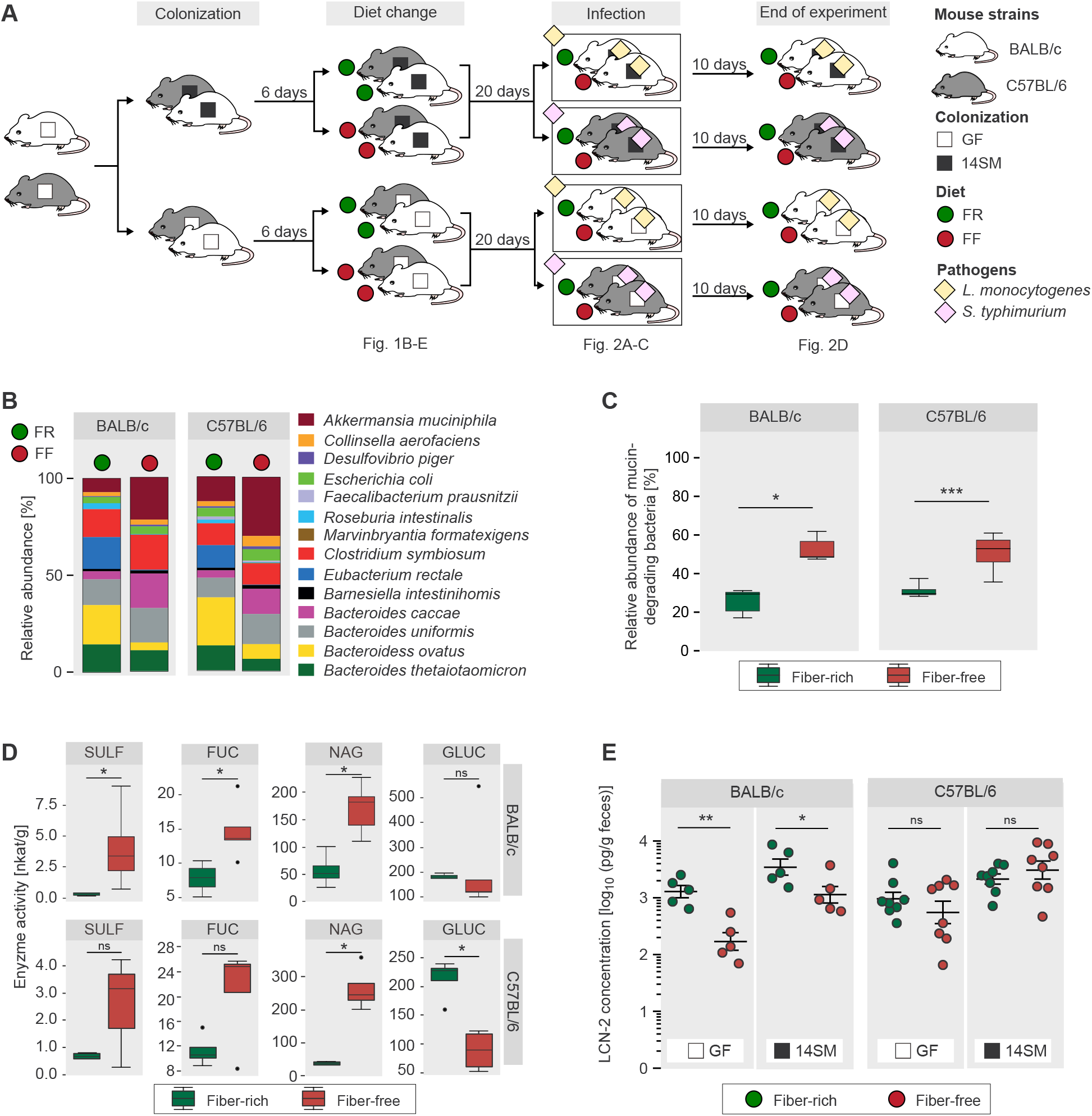
Fiber-deprivation increases abundance and activity of mucin-degrading gut bacteria in both BALB/c and C57BL/6 mice. (A) Experimental timeline. Half of the 6–10 weeks old, age matched GF BALB/c and C57BL/6 mice were gavaged with the 14SM gut microbiota on two consecutive days while the other half was maintained GF. Six days after the gavage, half of the mice from the GF and 14SM groups continued on the FR diet, while the other half were switched to the FF diet. The mice were maintained on their respective diets for 20 days and then BALB/c mice were infected with *L. monocytogenes* and C57BL/6 mice were infected with *S. typhimurium* after which the mice were observed for another 10 days. (B) Relative bacterial abundance before infection, determined by qPCR on DNA extracted from fecal pellets. While some low abundant bacteria might not be visible in the figure, the presence of all 14 bacteria was detected. (C) Combined relative abundances of four mucin-degrading bacteria *A. muciniphila*, *B. caccae*, *B. intestinihominis* and *B. thetaiotaomicron* using the same data from panel B. Tukey box plot, Mann–Whitney test. (D) Glycan-degrading enzyme activity of the gut microbiome determined by stool-based *p*-nitrophenyl glycoside-based enzyme assays. Sulfatase (SULF), α-fucosidase (FUC) and β-N-acetyl-glucosaminidase (NAG) are key mucindegrading enzymes, while β-glucosidase (GLUC) serves as a control for general glycan-degrading activity. Tukey box plot. Wilcoxon Rank Sum Test. (E) Fecal LCN-2 levels determined by ELISA on the day before the pathogen infection. Error bars represent SEM. Unpaired, two-tailed, t-test. BALB/c: n=5 mice/group. C57BL/6: GF FR group, n=7/group; other groups, n=8/group. Green: FR-fed mice; Red: FF-fed mice. ns, non-significant; \**p*<0.05; \*\**p*<0.01; \*\*\**p*<0.001; ****p<0.0001.

Throughout the feeding period before the pathogen infection, neither FR-nor FF-fed mice exhibited any obvious physiological abnormalities, irrespective of whether they were 14SM-colonized or not. In line with our previously published study with Swiss Webster mice hosting our 14SM community (4), fiber deprivation significantly shifted the gut microbiota of both BALB/c and C57BL/6 mice towards an increased relative abundance of the mucin-degrading bacteria *Akkermansia muciniphila* and *Bacteroides caccae* **(Fig. 1B)**. Whereas, the relative abundance of the typical fiber-degrading strains *Bacteroides ovatus*, *Eubacterium rectale* and *Roseburia intestinalis* decreased significantly in FF-fed mice of both genotypes compared to 14SM-colonized mice on the FR diet **(Fig. 1B)**. In contrast to *C. rodentium*, the primary infection site of these intracellular pathogens is in the small intestine and not the colon, yet previously it was shown that Muc2-deficiency renders colon as the main site for establishing a systemic spread of *L. monocytogenes* (8). Accordingly, the expansion of mucin-degrading commensals in FF-fed mice **(Fig. 1C)** prompted us to investigate the activity of bacterial mucin-glycan degrading enzymes, which would be a proxy for the erosion of the mucus barrier. We detected significantly increased fecal activities of key mucin glycan-degrading bacterial enzymes, such as sulfatase (SULF), α-fucosidase (FUC) and β-N-acetyl-glucosaminidase (NAG) in FF-fed BALB/c mice compared to FR-fed mice **(Fig. 1D)**. In C57BL/6 mice, only NAG was significantly increased while SULF and FUC showed a non-significant trend **(Fig. 1D)**. Moreover, the activity of β-glucosidase (GLU)—an enzyme that indicates microbial plant fiber metabolism—did exhibit significant changes in BALB/c mice and significantly decreased in FF-fed C57BL/6 mice **(Fig. 1D).** Overall, our results show an increased activity of the carbohydrate-active enzymes during fiber deprivation is specific to mucin glycan-degrading enzymes (**Fig. 1D)**.

These results indicate a diet-induced impairment of the colonic mucus layer, thereby increasing interactions between host cells and the intestinal microbiome. Thus, we determined potential diet-induced colonic inflammation via detection of fecal lipocalin-2 (LCN-2) levels, which is considered as a biomarker for low-grade inflammation (16). In BALB/c mice, we detected significantly increased levels of LCN-2 in both, FR-fed GF and FR-fed 14SM-colonized mice, compared to their FF-fed counterparts, while we did not detect any differences in C57BL/6 mice **(Fig. 1E)**. In contrast, in our previous study, 14SM-colonized Swiss-Webster mice show increased LCN-2 levels on the FF diet (4), suggesting that dietary fiber-mediated colonic baseline inflammation is likely dependent on the rodent genetic background. Furthermore, at least in BALB/c and C57BL/6 mice, this baseline inflammation is largely independent of the presence of the microbiota, as the observed trends were similar in GF mice **(Fig. 1E)**. Interestingly, we observe differences in the relative abundances of 14 strains in BALB/c, C57BL/6 **(Fig. 1B)** and Swiss Webster mice (4), despite being fed an identical FR diet, which indicates that the host genetic background plays a role in the colonization of our 14 strains.

After a 20-day feeding period, we infected both mouse strains with their respective pathogens **(Fig. 1A)**. Body weight and disease scores of all mouse groups were assessed daily for up to 10 days post infection (dpi). Lethality of *L. monocytogenes-infected* GF FR-fed BALB/c mice reached 100% by 4 dpi, while their FF-fed counterpart provided a significantly higher survival rate **(Fig. 2A, left panel)**. Similarly, 14SM-colonized FF-fed BALB/c mice had a significantly higher survival rate than their FR counterpart and intriguingly, all 14SM-colonized FF-fed animals survived the infection **(Fig. 2A, left panel)**. In accordance with previous reports stating that mice harboring an intestinal microbiota are less susceptible to *L. monocytogenes* infections than GF mice (11), 14SM-colonized BALB/c mice generally provided increased survival compared to the GF controls fed the same diet **(Fig. 2A, left panel)**. In line with the course of the survival curves, weight loss in FR-fed and *L. monocytogenes*-infected BALB/c mice, either 14SM-colonized or GF, was significantly higher compared to the corresponding FF-fed groups **(Fig. 2B, left panel)**. Additionally, daily-assessed disease scores in all four *L. monocytogenes*-infected BALB/c mouse groups (**Fig. 2C, left panel**; see **Table 1** for disease scoring scheme) underscore that susceptibility to *L. monocytogenes* infection is more dependent on the fiber content of the diet itself than on presence of a microbiota or its diet-influenced composition. Interestingly, fecal *L. monocytogenes* load did not significantly differ between both diets of the GF and 14SM groups, except for the last time point in the 14SM group which hints at a faster clearance in 14SM FR-fed mice (**Fig. 2D**). Systemic dissemination of *L. monocytogenes* in BALB/c mice was assessed by detection of CFUs in liver and spleen **(Fig. 2E)**. In contrast to the fecal pathogen levels, both FR-fed groups showed significantly increased dissemination of *L. monocytogenes* into liver compared to their FF-fed counterparts **(Fig. 2E, upper left panel)**. Similarly, dissemination into spleen was significantly higher in GF FR-fed mice compared to FF-fed controls **(Fig. 2E, lower left panel)**. These results suggest that the fiber-free diet does not affect growth of *L. monocytogenes*, but hinders its translocation across the epithelium.

**FIG 2.**
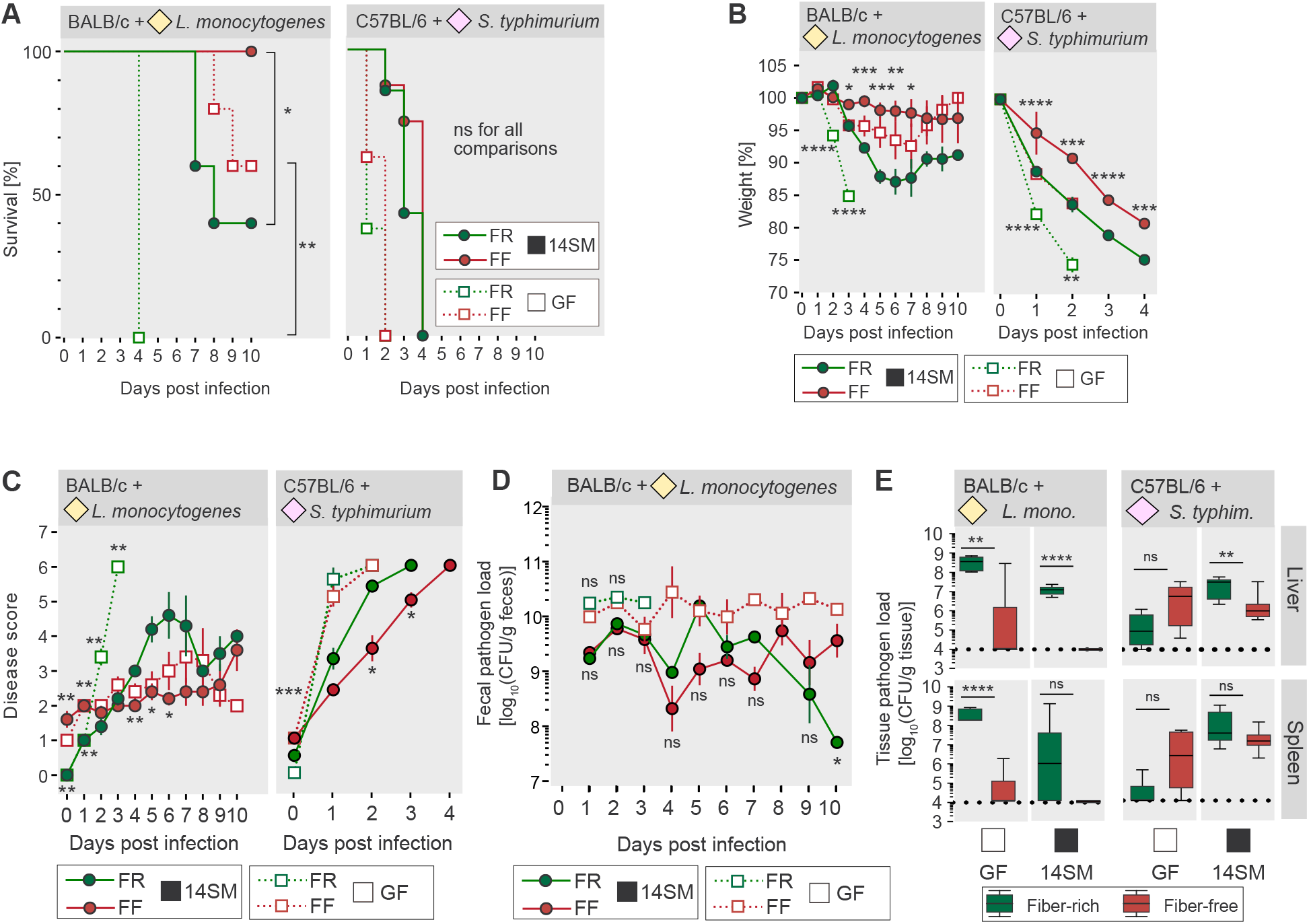
Fiber deprivation protects against *L. monocytogenes* and *S. typhimurium* in a microbiome-independent manner. (A) Survival curve of enteropathogen-infected mice. Log-rank test between both diets of GF or 14SM groups respectively. (B) Weight change of the enteropathogen-infected mice. Day 0 value was determined immediately before the gavage. Error bars represent SEM. Unpaired, two-tailed t-test between both diets of GF (bottom significance labels) or 14SM group (top significance labels); comparisons are not significant when the significance is not displayed. (C) Average disease score attributed to each enteropathogen-infected group. Day 0 value was determined immediately before the gavage. Error bars represent SEM. Mann–Whitney test between both diets of GF (top significance labels) or 14SM group (bottom significance labels); comparisons are not significant when the significance is not displayed. (D) Fecal *L. monocytogenes* load of BALB/c mice during the 10 days of infection. Depending on the sampling day, 1-5 samples per group were obtained and evaluated. Fecal *S. typhimurium* load in C57BL/6 mice could not be determined, as the mice did not consistently provide fecal material due to the severe disease. Tukey box plot; unpaired, two-tailed t-test. (E) Pathogen loads of liver and spleen tissues on the day each mouse was euthanized. Samples below the measurable threshold of 10^4^ CFU (dotted black line) were considered as 10^4^. Tukey box plot; unpaired, two-tailed t-test. BALB/c: n=5 mice/group. C57BL/6: GF FR group, n=7/group; other groups, n=8/group. Green, FR-fed mice; Red, FF-fed mice; unbroken lines, 14SM mice; dotted lines, GF mice. ns, non-significant; \**p*<0.05; \*\**p*<0.01; \*\*\**p*<0.001; ****p<0.0001.

**Table 1.** Scoring system used to determine disease severity of mice. Mice reaching the humane endpoint (HEP) were scored as maximum score of 6.

| Category | Score |
| --- | --- |
| **Body weight** |  |
| 5–10% weight loss | 1 |
| 11–15% weight loss | 2 |
| 16–20% weight loss | 3 |
| ≥20% weight loss | HEP |
| Pinched skin/dehydration | 4 |
| **Coat condition** |  |
| Coat slightly unkempt | 1 |
| Slight piloerection | 2 |
| Marked piloerection | 4 |
| **Body function** |  |
| Tachypnoea | 3 |
| Dyspnoea | 5 |
| **Environment** |  |
| Loose stools or diarrhoea | 1 |
| Blood in diarrhoea | HEP |
| **Behaviours** |  |
| Tense and nervous on handling | 3 |
| Markedly distressed on handling,<br>e.g. shaking, vocalizing, aggressive | 4 |
| **Locomotion** |  |
| Slightly abnormal gait/posture | 1 |
| Markedly abnormal gait/posture | 4 |
| Significant mobility problems<br>or reluctance to move | HEP |
| **Procedure-specific indicators** |  |
| Conjunctivitis | 4 |
| **Implementation of humane endpoint (HEP)** |  |
| Total score | ≥ 6 |

In contrast to *L. monocytogenes-infected* BALB/c mice, all *S. typhimurium-infected* C57BL/6 mice died within 4 days after infection **(Fig. 2A, right panel)**. Nevertheless, there were no significant differences in survival rates between FR-fed and FF-fed GF mice as well as between 14SM-colonized mice fed the two different diets **(Fig. 2A, right panel)**. Despite no significant differences in survival between *S. typhimurium-infected* C57BL/6 mice fed different diets, weight loss in FR-fed 14SM-colonized, as well as in FR-fed GF C57BL/6 mice, was significantly increased compared to their FF-fed 14-SM colonized or GF mice **(Fig. 2B, right panel)**. Disease scores of all mice reached the maximum possible score of 6, requiring immediate euthanasia, by 4 dpi **(Fig. 2C, right panel)**. Notably, GF mice were more susceptible to *S. typhimurium* infection than 14SM-colonized mice and FR-fed 14SM mice provided significantly higher disease scores than FF-fed 14SM-colonized mice. Furthermore, we detected no significant differences in dissemination of *S. typhimurium* into liver between both GF groups; in 14SM-colonized mice, FR-fed animals provided higher *S. typhimurium* CFUs in liver compared to FF-fed mice **(Fig. 2E, upper right panel)**. In spleen, however, no significant differences in CFUs were observed when comparing the diets of 14SM-colonized or GF mice **(Fig. 2E, lower right panel)**.

These results suggest that dietary fiber deprivation has a protective effect against the intracellular, food-borne pathogens *L. monocytogenes* and *S. typhimurium*, whose preferable infection site is the small intestine (12, 17). Our data suggest that this effect is microbiome independent and is more pronounced in BALB/c mice infected with *L. monocytogenes* compared to C57BL/6 mice infected with *S. typhimurium.* In contrast to Swiss Webster mice infected with cecum- and colon-targeting *C. rodentium* (4), elevated mucin degradation in BALB/c and C57BL/6 mice, as consequence of fiber deprivation, did not promote susceptibility to the chosen enteropathogens. Overall, we determined a direct impact of dietary fiber components on host susceptibility to enteropathogenic infections, which seems to be rooted in a heightened translocation efficiency. This cannot be counteracted by pathogenicity-modulating properties in 14SM-colonized mice, although the microbiota delayed the overall disease course and pathogen load. Our data suggest a potential pre-priming of the host in response to dietary fiber, which potentially facilitates subsequent pathogen infection. However, we cannot exclude the possibility that the fiber types present in our FR diets promote pathogen virulence, which cannot be counteracted by the 14SM microbiota. Indeed, a study in guinea pigs showed that supplementation with the dietary fibers pectin and inulin significantly increased the translocation of *L. monocytogenes* into liver and spleen (18). However, this study also shows that supplementation with galactooligosaccharides and xylooligosaccharides decreased the translocation (18), indicating a fiber-source specific virulence modulator of *L. monocytogenes.* In this context, increased fiber consumption has previously been linked to both, increased and decreased, susceptibility (13, 19, 20) to *S. typhimurium* infections, indicating that not only the presence or absence of dietary fiber in a mouse chow determines enteropathogen susceptibility, but the source or type of fiber is also an essential factor. Despite many advantages of gnotobiotic mouse studies (21), the potential absence of interactions between specific commensal bacteria and pathogens such as *Prevotella* spp. with *L. monocytogenes* (14) or *Mucispirillum shadleri* with *S. typhimurium* (22) must be considered as a limitation of our 14SM model. Moreover, gnotobiotic models might fail to provide a real-life picture of colonization resistance provided by a complex microbiome against both *L. monocytogenes* and *S. typhimurium* infections (11, 12). A potential caveat in comparing results obtained from our FR and FF diets could be the higher amount of simple sugar in the FF diet (4).

The intriguing impact of dietary fiber on increased susceptibility to enteropathogenic infections in mice, that is independent of the gut microbiota, calls attention to giving due importance when designing diets in mouse studies. Thus, mouse studies investigating underlying mechanisms of enteropathogen infections should involve a critical assessment of the animal chow composition across different laboratories. Our observation might have been overlooked in the absence of GF control groups, highlighting the importance of such controls when studying the enteropathogenesis mechanisms. At a broader level, our observational study suggests that potential dietary modulations via fiber supplementation for the benefit of human health should be performed carefully, considering the underlying microbiota composition and acknowledging potential downfalls due to unexpected side effects.

## Materials and Methods

### Ethical statement

All animal experiments were performed according to the *“Règlement Grand-Ducal du 11 janvier 2013 relatif à la protection des animaux utilisés à des fins scientifiques”* based on the Directive 2010/63/EU on the protection of animals used for scientific purposes and approved by the Animal Experimentation Ethics Committee of the University of Luxembourg and by the Luxembourgish Ministry of Agriculture, Viticulture and Rural Development (national authorization number: LUPA 2020/27). The mice were housed in isocages under gnotobiotic conditions in accordance with the recommendations stated by the Federation of European Laboratory Animal Science Association (FELASA).

### Experimental design and dietary treatment

Six to ten weeks old, age-matched male germ-free (GF) BALB/c (n=20, 5 per group) and C57BL/6N (n=31, GF Fiber-rich (FR) group: 7 per group, other groups: 8 per group) were housed in isocages with up to five animals per cage. Light cycles consisted of 12 hours of light and sterile water and diets were provided ad libitum. The GF status of the mice was confirmed by aerobic and anaerobic microbial culturing of fecal samples. As per the groupings, the relevant mice were gavaged with 0.2 ml of a 14-member synthetic human gut microbiota (14SM) gavage mix on two consecutive days. The gavage mix was prepared as described previously (15). Before and six days following the gavage, all mice were maintained on a standard mouse chow which we refer to as fiber-rich (FR) diet. Afterwards half of the gavaged and half of the GF mice were switched randomly to a fiber-free (FF) diet while the rest were maintained on the FR diet. In contrast to the FR diet, the FF diet does not contain dietary fiber from plant sources, but instead contains increased glucose levels (4). All mice were maintained for 20 days on their respective diet while fecal samples were collected once a week. After this 20-day feeding period, the BALB/c mice were infected with *L. monocytogenes* and the C57BL/6N mice were infected with *S. typhimurium*. Following the infection, the mice were observed for up to 10 days on their respective diets and fecal samples were collected daily for all possible mice. Upon reaching the humane endpoint or the end of the 10 days observation time, mice were euthanized by cervical dislocation. Liver and spleen were collected to determine pathogen load and spleen weight. Cecal contents were flash frozen and stored at −80 °C for LCN-2 level measurements (see below). Due to the rapid disease development, it was not possible to reliably obtain fecal material during the course of the *S. typhimurium* infection due to the severe symptoms, as such we could not compare CFU counts from feces between these groups.

### Animal diets

The fiber-rich diet was an autoclaved rodent chow (LabDiet, 5013), while the fiber-free diet was manufactured and irradiated by SAFE diets (Augy, France) according to the modified Harlan.TD08810 diet described previously (4).

### Colonization with 14-member synthetic microbiota (14SM)

All 14SM-constituent strains were cultured and intra-gastrically gavaged as described previously (15).

### Quantification of bacterial relative abundance

The colonization of individual strains in the 14-member synthetic microbiota was confirmed using phylotype-specific qPCR primers as described previously (15), and the relative abundances of individual microbial strains were computed using the same qPCR protocol (15).

### Pathogen culturing and enumeration

Both *Listeria monocytogenes (Murray et al.) Pirie (ATCC^®^ BAA-679™)* and *Salmonella enterica* subsp. *enterica* (ex Kauffmann and Edwards) Le Minor and Popoff serovar Typhimurium (Strain SL1344; DSM 24522) were grown overnight at 37 °C under aerobic conditions in Luria Bertani (LB) broth. Cultures were then spun down by centrifugation and resuspended in LB broth to reach the appropriate colony forming units (CFU) for gavage. BALB/c mice were infected with 10^9^ CFUs of *L. monocytogenes* and C57BL/6 mice were infected with 10^8^ CFUs of *S. typhimurium.* Fecal CFU enumeration was performed as described previously (4) with the modification of the selective media which differed based on the strain. Tissue was processed in the same manner, except that the homogenization was performed using a tissue grinder. *L. monocytogenes* was plated on Oxford agar plates, while *S. typhimurium* was plated on streptomycin-containing (50 μg/ml) LB agar plates.

### Mouse disease scoring

A project specific scoring system based on the FELASA guidelines for reporting clinical signs in laboratory animals (23) was used to determine mouse disease score. This scoring system is shown in **Table 1**.

### Lipocalin ELISA

Samples for the Lipocalin ELISA were prepared as described previously (4) and measured using the Mouse Lipocalin-2/NGAL DuoSet Elisa R&D Systems (Biotechne, Minneapolis, United States) according to the manufacturer’s instructions.

### Detection of bacterial glycan-degrading enzyme activities

Enzymatic activities of sulfatase, α-fucosidase, β-*N*-acetyl-glucosaminidase and β-glucosidase were determined using *p*-nitrophenyl glycoside-based enzyme assays from fecal samples as described previously (24).

## ACKNOWLEGMENTS

This work was supported by the Luxembourg National Research Fund (FNR) CORE grants (C15/BM/10318186 and C18/BM/12585940) to M.S.D.

We thank Pascale Cossart and Olivier Dussurget from the Pasteur Institute in Paris for their support and for providing us with the *Listeria monocytogenes* strain used in this work.

The authors declare no competing interests.

M.W., J.Z., and M.S.D. designed the study; M.W. and A.S. performed the experiments; M.W. and M.S.D. wrote the original manuscript draft; M.W., A.S., J.Z., and M.S.D. reviewed and edited the manuscript.

